# Nanoscale Organization of FasL on DNA Origami as a Versatile Platform to Tune Apoptosis Signaling in Cells

**DOI:** 10.1101/2020.07.05.187203

**Authors:** Ricarda M. L. Berger, Johann M. Weck, Simon M. Kempe, Tim Liedl, Joachim O. Rädler, Cornelia Monzel, Amelie Heuer-Jungemann

## Abstract

Nanoscale probes with fine-tunable properties are of key interest in cell biology and nanomedicine to elucidate and eventually control signaling processes in cells. A critical, still challenging issue is to conjugate these probes with molecules in a number- and spatially-controlled manner. Here, DNA origami-based nanoagents as nanometer precise scaffolds presenting Fas ligand (FasL) in well-defined arrangements to cells are reported. These nanoagents activate receptor molecules in the plasma membrane initiating apoptosis signaling in cells. Signaling for apoptosis depends sensitively on FasL geometry: fastest time-to-death kinetics are obtained for FasL nanoagents representing predicted structure models of hexagonal receptor ordering with 10 nm inter-molecular spacing. Slower kinetics are observed for one to two FasL on DNA origami or FasL coupled with higher flexibility. Nanoagents with FasL arranged in hexagons with small (5 nm) and large (30 nm) spacing impede signal transduction. Moreover, for predicted hexagonal FasL nanoagents, signaling efficiency is faster and 100× higher compared to naturally occurring soluble FasL. Incubation of the FasL-origami nanoagent in solution exhibited an EC50 value of only 90 pM. These studies present DNA origami as versatile signaling platforms to probe the significance of molecular number and nanoscale ordering for signal initiation in cells.

## Introduction

The astounding versatility and specificity by which biological signals are generated by cells are in part attributed to the cell’s ability to form characteristic molecular complexes in the plasma membrane. These complexes transduce a signal inside the cell, whenever an adequate stimulus is captured from the environment.^[1–4]^ Tools enabling acute control of such complex formation or its inhibition with effects on the signaling pathway are of primary interest in fundamental cell biology or nanomedicine.^[5]^ While functionalized nano-probes such as nanoparticles,^[6–8]^ quantum dots^[9–10]^ or nanodiscs^[11]^ already demonstrated to elucidate cell signaling behavior, the number and spatial localization of ligands on these probes are difficult to control. DNA origami, on the other hand, is a powerful tool to build versatile DNA-based platforms^[12–15]^ which satisfy multiple structural and biofunctional constraints including the possibility of complex molecular conjugation with biomolecules such as peptides or proteins with nanometric precision.^[16–23]^ Such defined molecular organization can provide unique insights into ligand-receptor interactions in signaling complexes and the resulting signal transduction as well as serve to enhance or block particular signals.^[24–25]^ Among the signaling complexes which present themselves as supramolecular assemblies, receptors and ligands of the tumor necrosis factor (TNF) super family are extensively studied and models of their multimerization and cluster formation have been proposed.^[26]^ Within this family, trimeric Fas ligand (FasL, CD178) plays a pivotal role in cell decision making towards proliferation or apoptosis.^[27–30]^ FasL is mainly expressed by lymphoid and myeloid cells and eliminates cancer cells by FasL-mediated apoptosis.^[31–34]^ However, recent studies demonstrated that tumor cells may also resist FasL-induced apoptosis, presumably due to Fas-mediated activation of other receptors^[35–36]^ or differences in the soluble and bound presentation of FasL.^[37]^ For these reasons, predominant interest exists in unraveling and modulating the key parameters leading to Fas receptor complex formation and unrestricted signaling. The present models of Fas signal initiation suggest that Fas receptor (FasR, CD95) may either pre-arrange in trimers assembling into hexameric patterns on the membrane (**Figure S1**) or remain as monomers and dimers.^[26, 38–39]^ Upon binding of trimeric FasL, up to three FasR may bind from different sites to the ligand. Depending on the ligand-receptor availability, they may further arrange in hexameric supramolecular structures with ~ 10 nm intermolecular spacing and recruit intracellular adaptor proteins to form the death-inducing signaling complex (DISC) eventually leading to apoptosis.^[26,29, 40–41]^

Nevertheless, the role of molecular ordering of FasL for signaling complex formation could not yet be deciphered experimentally, mostly due to a lack of tools enabling acute control over molecular positioning at the nanoscale. To address this question, we designed DNA-based nanoagents consisting of a one-layer DNA origami sheet functionalized with different nanometric arrangements of FasL to act as nanometric graded signal initiators. In brief, we constructed hexagonal and dimeric FasL patterns on DNA origami with varying FasL-FasL distances and probed the apoptotic kinetics of HeLa cells stably expressing FasR-mEGFP when exposed to these nanoagents (**Figure 1A**). Cell apoptosis rates were strongly affected by the size of FasL hexagons showing maximal efficiency at 10 nm intermolecular FasL-FasL distance and a five times reduced efficiency for smaller (5 nm) and larger (30 nm) patterns. Moreover, dose-response studies revealed a very low EC50 value of only 90 pM. Incubation of cells with the nanoagent enhanced cell apoptosis by 100× compared to free ligand in solution. Hence, FasL-DNA origamis can function as well-defined platforms to unravel effects of spatial molecular organization on the signal formation in cells.

**Figure 1.**
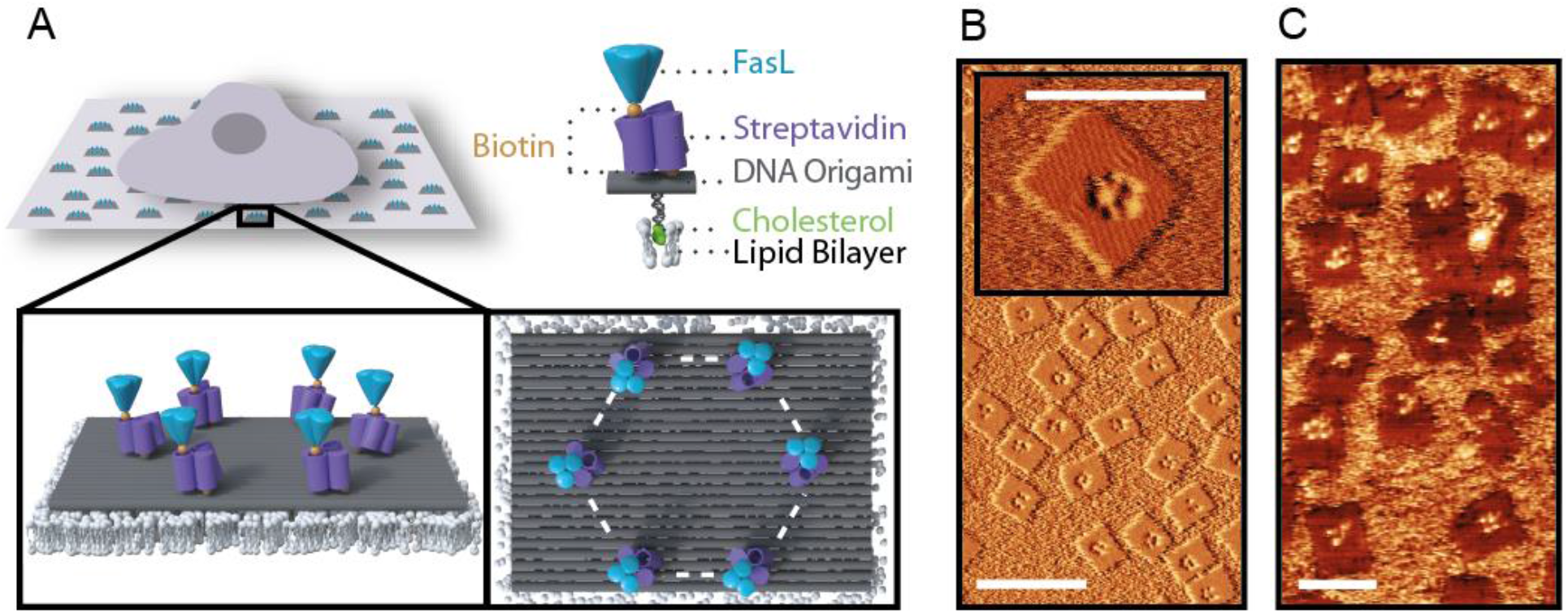
FasL functionalized DNA origami (FasL-nanoagent). (A) Schematic illustration of a cell seeded on DNA origami grafted to a supported lipid membrane. The magnifications show the Streptavidin-Biotin mediated linkage of trimeric Fas ligand to DNA origami. Top view illustrates the hexagonal arrangement indicated by white dashed lines. (B) AFM Lock-In Amplitude image of Streptavidin functionalized DNA origami sheets (200 nm scale bar). The insert exemplifies the hexagonal arrangement of Streptavidin (100 nm scale bar). (C) AFM image of DNA origami functionalized with Streptavidin and FasL (100 nm scale bar).

## Results and Discussion

The nanoagent used in this study is based on a rectangular DNA origami sheet consisting of 24 parallel interconnected helix bundles (dimensions: 100 × 70 nm), formed from a 7249 nt long circular scaffold and 200 staple strands (see **Figure S2**).^[14, 42]^ In order to achieve precise nanometric arrangement of FasL, DNA staple strands containing a terminal Biotin were designed to protrude from the DNA origami at designated positions resulting in various defined binding sites for Streptdavidin and subsequently FasL (see **Figure 1A**). The addition of Streptavidin allowed for the conjugation of FasL, pre-trimerized via a T4-FOLDON (see **Figure S3**) and containing a C-terminal Biotin, to the DNA origami sheet. Five different geometries were fabricated: three hexagonal arrangements with varying distances between adjacent FasL (5 nm, 10 nm and 30 nm) and two geometries with only two FasL placed at a distance of 10 nm and 20 nm, respectively (see **Figure S2B**). Atomic force microscopy (AFM) images show the DNA origami sheets adsorbed onto mica, displaying the designed hexagonal conformation of Streptavidins on each DNA origami (**Figure 1B**). We determined an average occupancy of 5 ± 0.3 Streptavidins per DNA origami, where 40 % ± 8 % of all analyzed DNA origamis were fully decorated with six Streptavidins (see **Figure 1B**). Subsequently, for DNA origamis decorated with Streptavidin and FasL, successful functionalization was confirmed by AFM imaging (**Figure 1C**) as well as by fluorescently labelled anti-FasL antibody staining (**Figure S4** and **Supplementary Movies M1**). Analysis of the AFM data revealed a 76 % ± 9 % binding efficiency of FasL to Streptavidin anchors with a positioning accuracy of ~ 5 nm estimated from the protein structures. While the positioning of Biotins on the DNA origami is nanometer precise, spatial positioning of the FasL is limited by the size (~5 nm) and multivalency of wild type (wt) Streptavidin.^[43]^ Mulivalency also imposes uncertainty on the actual number of bound ligands. To address this issue, we performed experiments with monovalent (mv) Streptavidin and found the same qualitative behavior as for the multivalent Streptavidin assemblies.

Having successfully established spatially designed, multivalent nanoagents, we investigated the cell apoptosis response when exposed to different FasL-DNA origamis, either presented on a supported lipid membrane (SLM) or in solution. In order to anchor FasL-DNA origamis to the SLM, a set of 8 staple strands at the bottom side of the DNA origami were modified with cholesterol moieties, capable of inserting into the SLM as previously reported.^[44–46]^ FasL-DNA origamis adsorbed on the SLM remained oriented and laterally mobile, hence perfectly addressable to cells. For the apoptosis assay, HeLa cells overexpressing FasR were exposed to the nanoagent-functionalized SLMs and analyzed via time-lapse imaging. First assays were carried out using FasL-DNA origamis with a hexagonal pattern and FasL-FasL distances of 10 nm, which are predicted to be an optimal distance for apoptosis induction.^[26, 47]^ Cells attaching to FasL-DNA origamis were observed to spread within the first three hours and subsequently exhibited pronounced blebbing and rounding-up, indicative of apoptosis (**Figure 2A** upper row**, Figure S5** and **Supplementary Movies M2**). In contrast, cells deposited on blank DNA origamis grafted to SLMs showed long-term normal spreading and cell division, indicating no negative effect of the bare DNA origamis on cells (**Figure 2A** lower row). Origami-free controls of cells, where FasL was bound to SLMs directly (via cholesterol-DNA-Streptavidin linkers) and distributed homogeneously, as well as cells on bare SLMs in the presence of soluble FasL showed apoptosis over distinctly slower time scales and for a lower number of cells than FasL-DNA origamis with hexagonal FasL clusters. **Figures 2B** and **C** show the corresponding cell-death kinetic curves and boxplots. Here, the time course of the normalized cumulative sum of apoptotic events are compared for FasL-DNA origami, origami-free FasL bound to SLMs or FasL in solution. While for concentrations of 0.1 nM and 1 nM of FasL-DNA origami no significant change in the time-to-death kinetics is observed, apoptosis times for origami-free soluble and membrane-bound FasL are concentration dependent. Most importantly, a higher percentage of cells die within a short time span when subjected to FasL-DNA origamis compared to cells exposed to membrane-bound FasL or FasL in solution, which indicates that signal induction is dominated by the structural pre-arrangement of FasL. To quantitatively compare the potency of apoptosis induction of the various FasL presentations, the time-to-death (τ_*i*_) of all individual time courses was evaluated. We define the inverse of the individual time-to-death (τ_*i*_) value as apoptosis rate 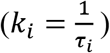. **Figure 2C** shows box plots of the apoptosis rates derived from a minimum of 500 cells per condition and with at least two repetitive experiments per condition.

**Figure 2.**
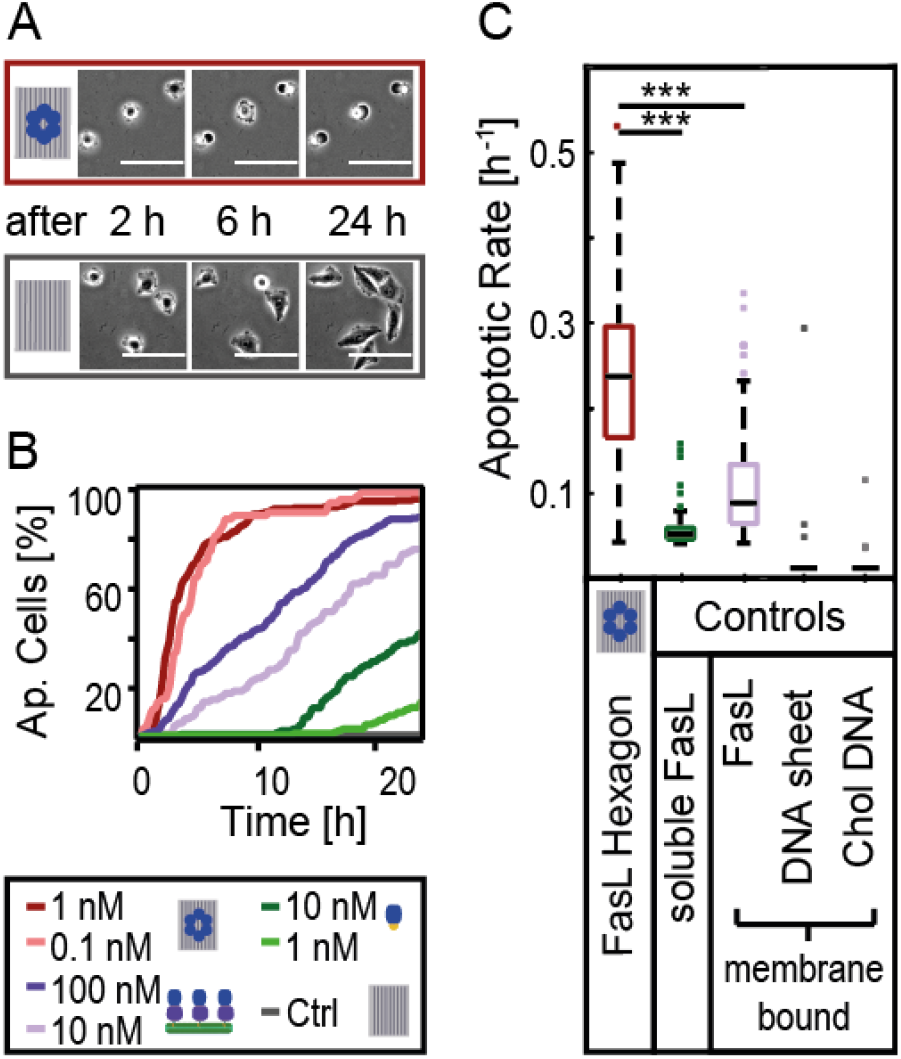
FasL induced apoptosis. (A) Comparison of representative bright field images showing morphological changes of cells seeded on FasL-nanoagent (red) or blank DNA origami (grey) on a supported lipid membrane after 2 h, 6 h, and 24 h. (B) Cell death kinetics from single experiments with at least 500 analyzed cells. FasL induced apoptosis is most efficient for the nanoagent (red) compared to FasL functionalized lipid membrane (lilac) and FasL in solution (green). (C) Efficiency of apoptosis initiation of nanoagent compared to origami-free FasL. Median of apoptosis rate for 1 nM nanoagent is significantly higher than soluble FasL (1 nM) or membrane-bound FasL (10 nM). DNA origami sheet or Cholesterol coupled DNA do not affect cells. n > 500 cells/condition. Ranksum test: ***p < 0.001, **p < 0.01, *p < 0.05.

The median of the apoptosis rates is a measure of the efficiency of apoptosis induction and turns out to be higher for FasL-DNA origamis compared to origami-free FasL, i.e., membrane-bound FasL or FasL in solution. Here, this high efficiency was obtained despite the fact that membrane-bound FasL and FasL in solution are present at 10× higher concentrations compared to FasL-DNA origamis. These results are in agreement with findings by Holler *et al*., who fused two trimeric FasLs to form FasL-dimers based on immunoglobulins that showed elevated potency to induce apoptotic signaling.^[48]^ In addition, Zhang *et al.* used a Fas peptide to target FasR and to trigger apoptosis signaling.^[49]^ In their case, Fas peptide was coupled to a DNA tetrahedron to increase its local concentration, which resulted in an eight-fold higher potency relative to the soluble peptide (5 μM vs 40 μM Fas peptide concentration). Note, that in comparison to their study, our FasL-nanoagents displayed a significantly higher potency while the concentration of nanoagent was four to five orders of magnitude lower (0.1 nM).^[49]^ Overall, these results demonstrate that oriented and spatially pre-clustered FasL exhibit maximal efficiency to induce cell death signaling.

Having established the excellent potency of FasL-DNA origamis in comparison to membrane-bound or solution FasL, we next performed a series of proof-of-concept experiments to uncover how the nanometric arrangement of FasL affects the signaling initiation for apoptosis. To this end, three sizes of hexagonal FasL arrangements on DNA origami were designed exhibiting 5 nm, 10 nm or 30 nm intermolecular FasL-FasL separation. Intriguingly, **Figure 3A** (upper left panel) shows that 10 nm FasL spacing induces cell death significantly more efficiently (5× faster) than 5 nm or 30 nm spacing (see **Supplementary Table S6** for peak locations of Gaussian fit distributions to the histograms). These findings are in very good agreement with models of protein complex formation of the TNF superfamily, to which FasR and FasL belong.^[41, 47]^ In a recent review by Vanamee *et al.*, for example, the intermolecular distance between ligand-receptor pairs of the TNF superfamily was summed up to amount to ~ 12 nm.^[26]^ In the work of Gülcüler Balta *et al.* an optimal average FasL-FasL distance of about 9-11 nm for membrane anchored FasL was reported.^[47]^ However, in the latter study FasL was free to diffuse laterally, whereas in the present work the spatial arrangement of DNA origami-bound FasL was fixed.

**Figure 3.**
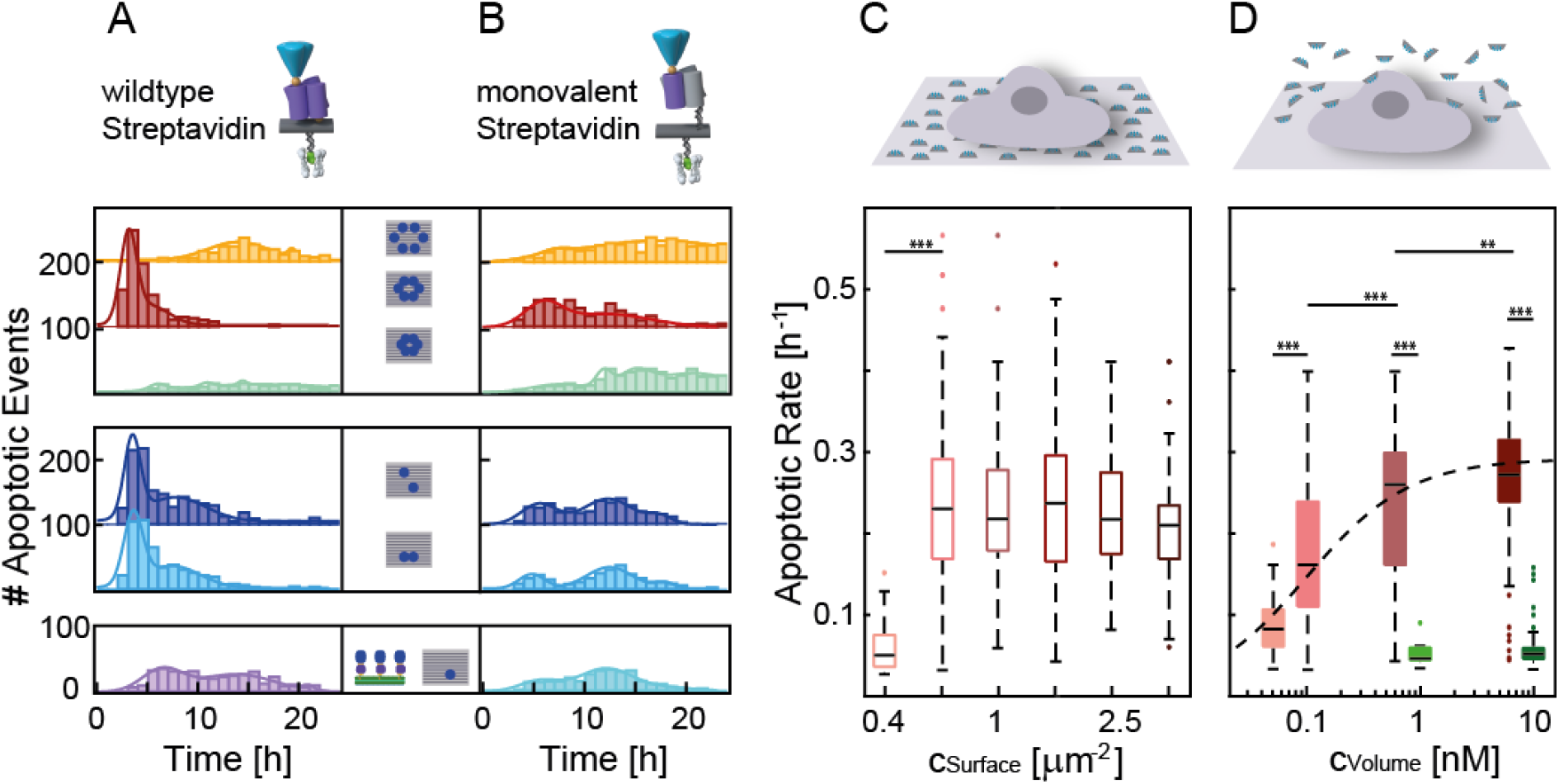
Cell death kinetics as a function of FasL distance, functionalization method and concentration. (A) Histogram of time-to-death data for different FasL-FasL distances arranged on DNA origami via wt Streptavidin: FasL hexagons with 10 nm intermolecular distance (red) exhibit highest efficiency in triggering cell apoptosis. FasL hexagons of 30 nm (orange) or 5 nm (mint) intermolecular distance, are least efficient. Two FasL arranged on the DNA origami with an inter-ligand distance of 20 nm (dark blue) or 10 nm distance (light blue) induce apoptosis faster compared to 100 nM membrane bound FasL (purple). (B) Same experiment as (A) but employing mv Streptavidin as linker. Additionally, data of a single FasL on DNA origami (cyan) is shown. The relative change between experiments follows the trend observed in (A) with broader distributions and shifted to longer times. Histogram data was fitted with multipeak Gaussian functions. (n > 500 cells/condition). (C) Boxplot with median of apoptotic rate for hexagonal FasL-nanoagent (10 nm spacing) at different surface concentrations of DNA origamis. The surface concentration was calculated from single molecule fluorescence analyses (SI). (n > 100 cells/condition). (D) Apoptotic rate as a function of volume concentration of soluble FasL hexagon nanoagent (10 nm, red) compared to soluble FasL (green). Fit: 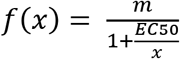 with m = 0.29 ± 0.02 and *EC*50 = 0.09 ± 0.02 nM, see text for further information (n > 100 cells/condition). Ranksum test: ***p < 0.001, **p < 0.01, *p < 0.05.

To decipher more precisely the effect of FasL number versus their arrangement, we fabricated two nanoagents displaying only two FasL molecules with an inter-ligand distance of either 10 nm or 20 nm. Remarkably, reducing the number of FasL molecules per DNA origami by a factor of three appeared to have only a moderate effect on the average time-to-death (**Figure 3A** middle left panel) in comparison to different FasL hexagon spacings, suggesting that already a low number of FasL can effectively induce apoptosis. A Ranksum test yielded a highly significant deviation (p < 0.001) between the distributions of hexagonal FasL-DNA origami and FasL-DNA origami with only two ligands, whereas a less significant deviation (p < 0.01) was found for FasL-DNA origami displaying two FasL at inter-ligand distances of 10 nm and 20 nm, respectively. On average, DNA origamis displaying two FasL at 10 nm inter-ligand distance were more potent than those with 20 nm inter-ligand distance. Yet, both were significantly more effective than origami-free, membrane-bound FasL (**Figure 3A** lower left panel). Intriguingly, histograms of FasL nanoagents displaying two FasL as well as the histogram of membrane-bound FasL exhibited multiple peaks indicating populations of cells with different apoptotic rates. To understand the origin of these peaks, we studied the time-to-death kinetics of HeLa cells as a function of FasR expression (see **Figure S7**). The variation in FasR expression was narrow as expected for a sorted cell line and no correlation between FasR expression and time-to-death values was found. By similar analyses, we could exclude that cell division influenced apoptosis kinetics (see **Figure S10**). Another explanation points to a diffusive rearrangement process, where bound FasL lateral rearrangement on the SLM or in solution requires additional time until a signal initiation complex is formed. Finally, an occurrence of multiple apoptotic timings was also observed by Márquez-Juarado *et al.* for TRAIL-induced apoptosis and was attributed to variability of mitochondrial expression in the cell population.^[50]^ Such variable mitochondrial expression levels may also occur in our cell line, even though we would expect its effects to appear in all histograms and not predominantly in those data, where FasL is mobile. In total, our results confirm that apoptosis initiation sensitively depends on FasL arrangements, where intermolecular distance has a predominant effect compared to FasL number. Thus, pre-formed FasL clusters with the correct distance may facilitate the formation of a hexameric pattern of the DISC,^[26, 34, 38–41]^ whereas pre-arranged FasL patterns with spacings that do not match the predicted optimal FasL-FasL distance (e.g. 5 nm or 30 nm) suppress Fas-signaling possibly by spatial mismatch. Similar phenomena were also reported in the field of immuno-activation, where artificial arrays of antigens with defined spacing showed an enhanced immune response of B cells.^[51]^

To probe effects of FasL linker flexibility and coupling, we then repeated the above experiments using recombinant mv Streptavidin instead of wt Streptavidin (**Figure 3B** right panels).^[52–54]^ Here, mv Streptavidin has only one single functional binding pocket for Biotin and a thiol-maleimide coupling to a defined handle on the DNA origami with a 20 bp linker enabled FasL spatial pre-orientation with 10 nm spatial flexibility (20 bp × 0.34 nm = 6.8 nm for DNA coupling in addition to the size of the Streptavidin), (see **Experimental Section**). Histograms of the corresponding time-to-death data revealed that in this case, apoptosis kinetics are significantly slower compared to that of the nanoagents formed with wt Streptavidin. However, the key observations for FasL-DNA origamis formed with wt Streptavidin are well reproduced: the 10 nm hexagon arrangement of FasL still appears as the most potent inducer of apoptosis, while for smaller and larger hexagonal arrangements apoptosis induction is suppressed. FasL-DNA origami displaying two ligands also resulted in slightly faster apoptosis rates than DNA origamis displaying only one FasL (**Figure 3B** lower right panel). Hence, mv Streptavidin coupling confirmed the general trend detected for the wt Streptavidin coupling, but the increasing linker flexibility and lack of precise ligand pre-orientation inhibits efficient apoptosis signal induction.

Having identified the most potent FasL configuration, we then probed how the cell signaling response could be optimized as a function of FasL-DNA origami concentration and performed a dose-response analysis. We exposed cells to decreasing amounts of FasL-DNA origami (hexagonal arrangement with 10 nm inter-ligand spacing) and calculated the surface concentration according to independent calibration experiments (see **Figure S9**). Intriguingly, we found a threshold-like behavior, where enhanced apoptosis rates as a function of surface concentration of DNA origamis are observed above 0.6 DNA origamis per μm^2^ with no further increase between 0.6 and 4 DNA origamis per μm^2^. Below 0.6 DNA origamis per μm^2^ the apoptosis rate drops significantly from 0.21 h^−1^ ± 0.01 h^−1^ to 0.05 h^−1^ ± 0.02 h^−1^ (**Figure 3C** and **Figure S10A**). These dose-response characteristics suggest that the apoptotic potency is saturated above a critical surface concentration of *c*_*Surface*_ = 0.6 DNA origamis per μm^2^. For a typical cell area of 1600 μm^2^ this corresponds to about 100 DNA origamis per cell to efficiently induce apoptosis.^[55]^

Finally, and in view of nanoagent applications in cell biology and nanomedicine, it is important to probe the potency of nanoagents for signal induction in comparison to the naturally occurring free ligand in solution rather than bound to the SLM. Thus, we seeded cells on the SLM and incubated them with hexagonal FasL-nanoagents (10 nm inter-ligand spacing, no cholesterol anchors) in the medium. **Figure 3D** and **Figure S10B** show how the apoptosis rate increases continuously with increasing volume concentration and how it asymptotically reaches saturation. Fitting the data, we thus obtain an EC50 value of 0.09 nM ± 0.02 nM, which is clearly lower than an EC50 value of 10 nM reported for soluble FasL^[56]^ and comparable to or even lower than values reported for other FasR targeting nanoagents.^[48–49, 57]^ Moreover, comparing FasL-DNA origamis incubated in solution with the cell response after incubating origami-free, soluble FasL we find that FasL-DNA origami with hexagonal arrangement and 10 nm inter-ligand spacing exhibit a 6× higher apoptotic rate of 0.27 h^−1^ ± 0.04 h^−1^ than soluble FasL (0.05 h^−1^ ± 0.01 h^−1^) even when the latter was applied at a significantly higher volume concentration of 10 nM.

## Conclusion

To conclude, we successfully engineered spatially pre-clustered FasL-DNA origami nanoagents as signaling platforms for cell death initiation. For the first time, systematic tuning of FasL nanometric spacing, their lateral organization, and linker flexibility enabled to identify benchmarks in cell signal initiation and sensitive changes in the corresponding cellular response. In particular, hexagonal FasL arrangements on DNA origami with 10 nm intermolecular spacing exhibited maximal potency as apoptosis inducer thereby confirming structural models of FasL signaling complex formation. Deviations from this intermolecular spacing lead to pronounced suppression of apoptosis signaling. Moreover, FasL-DNA origami required a 100× lower molar concentration to initiate apoptosis compared to the naturally occurring FasL molecules in solution and hence exhibited significantly higher potency. This constitutes a valuable result for future nanomedical approaches. We envision broad applicability of these nanoagents to decipher the physicochemical mechanisms underlying signal formation in cells, as for example the existence of local concentration thresholds or effects of molecular flexibility. Nanoagents may also provide unique insights into the structural organization of signaling complexes in cells as well as serve as efficient trigger of cluster-mediated cell responses in biological or medical applications.

## Experimental Section

### DNA origami fabrication

For the design of the DNA origami one layer sheet caDNAno 2.0 and the Picasso Software were used. ^[42, 58]^ The scaffold p7249 was purchased from tilibit nanosystems GmbH (Garching, Germany) and the staple oligonucleotides with the caDNAno generated sequences from Eurofins Genomic (Ebersberg, Germany). Also, biotinylated and fluorescently labelled oligonucleotides were produced by Eurofins Genomic. Cholesterol-TEG modified oligonucleotides were purchased from Biomers (Ulm, Germany). 10 nM scaffold was mixed with a 10× excess of staple oligonucleotides except fluorescent oligos (30× excess) and biotinylated oligonucleotides (80× excess) in a 1× TAE and 12.5 mM MgCl2 buffer. The complementary handle sequences for the Cholesterol oligonucleotides were added with a final concentration of 8 nM. DNA origami structures were annealed in 16 h temperature ramp from 65 °C to 20 °C. Correct folding was screened either with gel electrophoresis or AFM. Unpurified structures were stored at 4 °C until further use.

### AFM

First, DNA origami structures were purified 5× with Amicon^®^ Ultra-0.5 Centrifugal Filters, purchased from Merck (Darmstadt, Germany). Then, DNA origami structures were incubated on a mica at 1.5 nM concentration for 10 min. Both liquid and dry imaging were carried out on a Nanowizard Ultra Speed2 (Bruker Nano GmbH, Berlin, Germany). For liquid measurements 1.5 mL 11 mM MgCl_2_ (origami buffer) was added prior to imaging with tapping mode using either a FASTSCAN-B tip (Bruker Nano GmbH, Berlin, Germany) or a BL-AC40 TS tip (Oxford Instruments Asylum Research, Goleta CA, USA). For **Figure 1C** the sample was not purified after FasL addition resulting in adsorbed FasL surrounding the DNA origami sheets. For dry measurements samples were washed 3× with water, then blow-dried with Nitrogen and a OMCL-AC160TS tip from Olympus Corporation (Shinjuko, Japan) was used.

### Protein

Streptavidin was bought from Merck KGaA (Darmstadt, Germany) and the biotinylated FasL trimer was purchased from Apogenix (Heidelberg, Germany) and stored aliquoted and sterile at −20 °C. mv Streptavidin was produced as described by Sedlak *et al.* and stored in phosphate buffered silane (PBS) at 4 °C until further use.^[54]^ For thiol-maleimide coupling the mv Streptavidin was reduced with 1 mM TCEP over 30 min. Afterwards it was desalted with Zeba spin desalting columns (ThermoFisherScientific, Waltham MA, USA) and transferred into coupling buffer (50 mM Na_2_HPO_4_, 50 mM NaCl, 10 mM EDTA, pH 7.2). 10× maleimide-DNA (Biomers, Germany) was added and incubated for 60 min. Access maleimide and coupling buffer were removed with Amicon® Ultra-0.5 Centrifugal Filters and the functionalized mv Streptavidin stored in PBS at 4 °C until further use. In this form, the mv Streptavidin exhibiting a cysteine-side chain could then hybridize to complementary handle sequences protruding from the DNA origami.

### Surface functionalization

18:1 (Δ9-cis) DOPC was purchased from Avanti Polar Lipids, Alabaster, USA dissolved in chloroform solution. One-day prior measuring, one milligram of the lipids was pipetted with a glass syringe from Hamilton (Reno, USA) to a chloroform cleaned glass vial. Afterwards chloroform was evaporated by nitrogen gas and dried overnight under vacuum. Under sterile working conditions, precision cover glasses No. 1.5H (Paul Marienfeld GmbH & Co. KG, Lauda-Königshofen, Germany) were cleaned with UV grade Isopropanol and lens cleaning tissues, and were glued to ibidi stricky-Slide VI 0.4 (ibidi, Martinsried, Germany) and also stored overnight under vacuum. On the next day, 1 mL PBS was added to the dried lipids. The solution was vortexed, and tip sonicated on ice for at least 45 min until a clear solution was observable. In between, the slides were washed 5× with PBS. Then, 100 μl lipid solution was added to each channel and incubated for 1 h at room temperature. Afterwards, channels were washed twice with sterile DI water to evoke an osmotic shock. Next the lipid bilayer was washed 5× with PBS before adding Cholesterol DNA (100 nM). After incubating for 15 min, channels were washed twice with PBS and twice with origami buffer. Next, unpurified DNA origami structures were incubated for 30 min to hybridize to the Cholesterol DNA on the membrane. Then, channels were washed twice with origami buffer, twice with buffer A (10 mM Tris-HCL, 100 mM NaCl, pH 8.0) and Streptavidin was added at a concentration of 2.8 μM. After 10 min, the channels were rinsed with buffer A and FasL was incubated for 10 min at a concentration of 170 nM and subsequently washed with buffer A and with L15 (ThermoFisherScientific, Waltham MA, USA) medium with 10 % fetal bovine serum (FBS).

### Cell Culture and time-lapse measurements

Hela cells were cultivated in DMEM Glutamax (ThermoFisherScientific, Waltham MA, USA) and 10 % FBS. For time-lapse measurements medium was changed to L15 and 10 % FBS. 2000 cells were seeded per channel and Anti-Evaporation Oil (ibidi, Martinsried, Germany) was added on top of the channel. Immediately, the slides were transferred to the microscope chamber preheated to 37 °C. Alternatingly, brightfield and fluorescence images were recorded every 10 min for 20-30 h.

### Analysis

Time-lapse movies were analyzed with the ImageJ Timeseries analyzer plugin v3, where apoptotic blebbing was determined for each cell. Afterwards, the event times were processed with MATLAB (Mathworks), where a code was written for the statistical analysis.

## Supporting Information

Supporting Information is available.

## Acknowledgements

R.B. was supported by a Deutsche Forschungsgemeinschaft (DFG) Fellowship through the Graduate School of Quantitative Biosciences Munich (QBM).

T.L., J.R. and A.H-J acknowledge the financial support of the Deutsche Forschungsgemeinschaft DFG through SFB1032 “Nanoagents” (A06 and A07). CM acknowledges financial support of the Deutsche Forschungsgemeinschaft DFG through SFB1208 “Identity and Dynamics of Membrane Systems” (A12) and Volkswagen Foundation.

We thank S. Sedlak for providing mSA for our measurements and Joél Beaudouin (Institut de Biologie Structurale IBS, Grenoble) for providing plasmids and cell lines.

---

**Table of Contents Figure.**
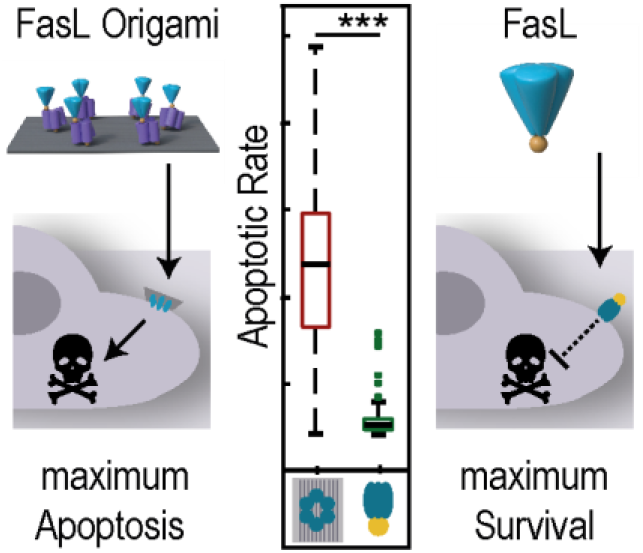
DNA origami allows for nanometrically-controlled arrangement of Fas ligands, which are heavily involved in apoptosis signaling. FasL-DNA origami nanoagents displaying a hexagonal ligand arrangement with inter-ligand distances of 10 nm exhibit a much faster and 100 × higher signaling efficiency in HeLa cells than soluble FasL without pre-arrangement.

