## Supplementary Information for "Nanoscale Organization of FasL on DNA Origami as a Versatile Platform to Tune Apoptosis Signaling in Cells"

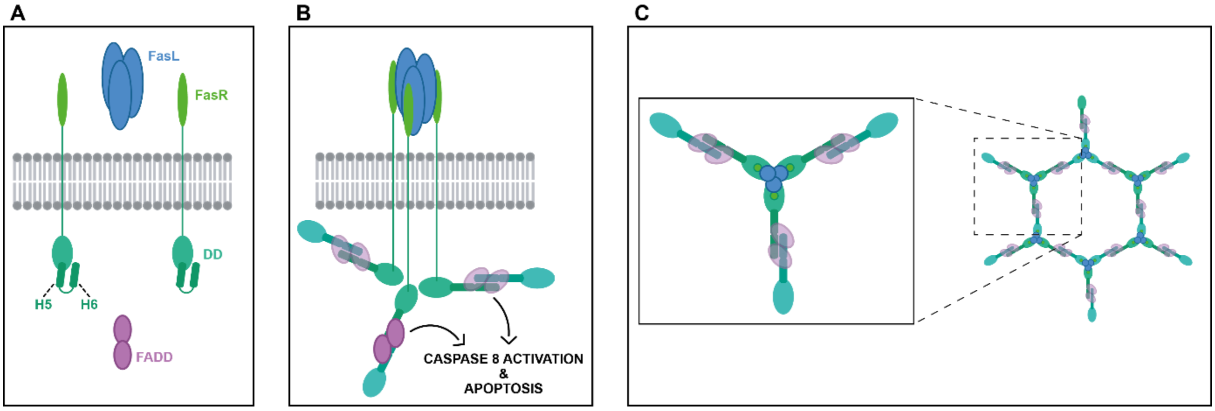


**Figure S1** Schematic illustration of the FasL induced apoptosis signaling pathway in cells. The here described model follows the description of Vanamee et al. [1] (A) When FasL is not bound to FasR, putatively the intracellular domain of FasR is in an inactive conformation. (B) Upon binding of FasL a conformational change in the death domain (DD) of FasR may allow Fas-associated-death domain (FADD) recruitment and coupling of DDs of neighboring FasR. Successive coupling between neighboring trimeric FasL-FasR is suggested to result in supramolecular complexes of a hexagonal structure. Autocatalysis of (pro-)caspases bound to the intracellular part of this hexagonal complex then initiate cell apoptosis. (C) Top view of hexameric cluster formation upon FasL binding with an expected intermolecular spacing between 9 – 15 nm according to literature.[1-3]

A

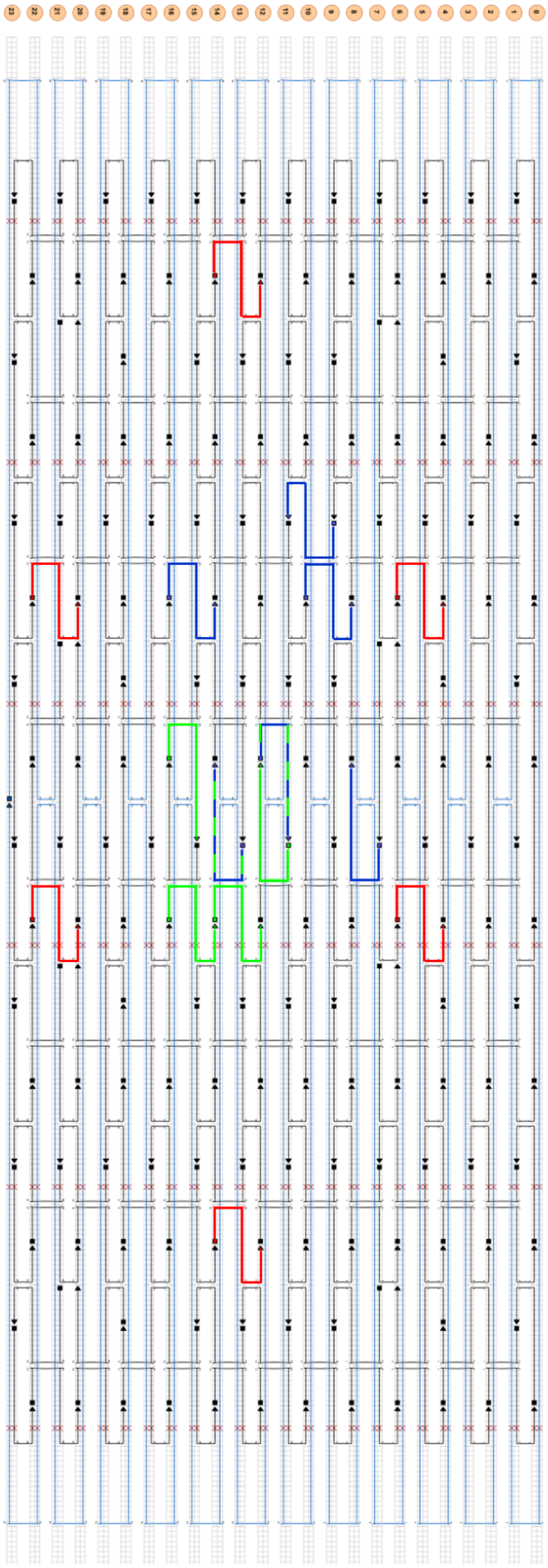

B

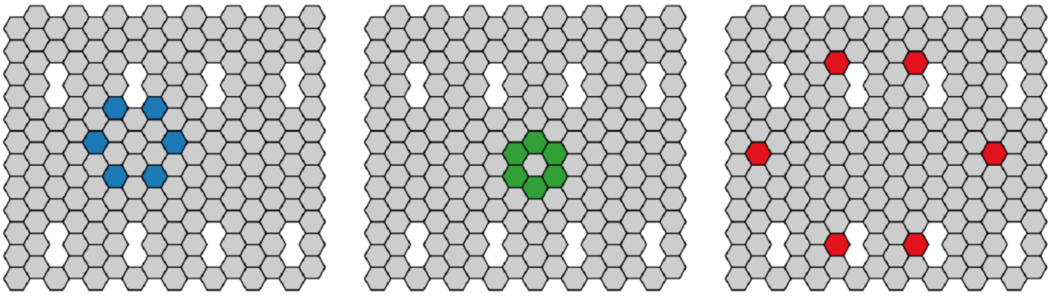

C

| Functionalized strands | Pattern | Position | Sequence |
|---|---|---|---|
| Cholesterol Handles | all | 4[127] | CATTCTCCTATTACTACCTTGTGTCGTGACGAGAAACACCAAA TTTCAACTTTAAT |
| | | 18[127] | CATTCTCCTATTACTACCGCGATCGGCAATTCCACACAACAGG TGCCTAATGAGTG |
| | | 18[191] | CATTCTCCTATTACTACCCACCCTCAGAAACCATCGATAGCAT TGAGCCATTTGGGAA |
| | | 18[255] | CATTCTCCTATTACTACCAACAATAACGTAAAACAGAAATAAAA ATCCTTTGCCCGAA |
| | | 18[63] | CATTCTCCTATTACTACCATTAAGTTTACCGAGCTCGAATTCG GGAAACCTGTCGTGC |
| | | 4[191] | CATTCTCCTATTACTACCCACCCTCAGAAACCATCGATAGCAT TGAGCCATTTGGGAA |
| | | 4[63] | CATTCTCCTATTACTACCATAAGGGAACCGGATATTCATTACG TCAGGACGTTGGGAA |
| | | 4[255] | CATTCTCCTATTACTACCAGCCACCACTGTAGCGCGTTTTCAA GGGAGGGAAGGTAAA |
| Cholesterol | Membrane anchor | | GGTAGTAATAGGAGAATG-Cholesterol TEG |
| FasL | 5 nm Hexagon | 13[160] | GTAATAAGTTAGGCAGAGGCATTTATGATATTTT[BIO] |
| | | 11[160] | CCAATAGCTCATCGTAGGAATCATGGCATCAATT[BIO] |
| | | 16[143] | GCCATCAAGCTCATTTTTTAACCACAAATCCATT[BIO] |
| | | 12[143] | TTCTACTACGCGAGCTGAAAAGGTTACCGCGCTT[BIO] |
| | | 16[175] | TATAACTAACAAAGAACGCGAGAACGCCAATT[BIO] |
| | | 14[175] | CATGTAATAGAATATAAAGTACCAAGCCGTTT[BIO] |
| | 10 nm Hexagon | 8[144] | TTATTACGAAGAACTGGCATGATTGCGAGAGGTT[BIO] |
| | | 11[95] | CGAAAGACTTTGATAAGAGGTCATATTTCGCATT[BIO] |
| | | 14[112] | TGTAGCCATTAAAATTCGCATTAAATGCCGGATT[BIO] |
| | | 11[159] | TTCTACTACGCGAGCTGAAAAGGTTACCGCGCTT[BIO] |
| | | 14[144] | GTAATAAGTTAGGCAGAGGCATTTATGATATTTT[BIO] |
| | | 8[112] | TTGCTCCTTTCAAATATCGCGTTTGAGGGGGTTT[BIO] |
| | 30 nm Hexagon | 14[47] | AACAAGAGGGATAAAAATTTTTAGCATAAAGCTT[BIO] |
| | | 22[111] | GCCCGAGAGTCCACGCTGGTTTGCAGCTAACTTT[BIO] |
| | | 6[111] | ATTACCTTTGAATAAGGCTTGCCCAAATCCGCTT[BIO] |
| | | 22[175] | ACCTTGCTTGGTCAGTTGGCAAAGAGCGGATT[BIO] |
| | | 6[175] | CAGCAAAAGGAAACGTCACCAATGAGCCGCTT[BIO] |
| | | 14[239] | AGTATAAAGTTCAGCTAATGCAGATGTCTTTCTT[BIO] |
| FasL(Maleimide) | 5 nm Hexagon | 13[160] | GTAATAAGTTAGGCAGAGGCATTTATGATATTTTTTCATTCTC TATTACTACC |
| | | 11[160] | CCAATAGCTCATCGTAGGAATCATGGCATCAATTTCATTCTCC TATTACTACC |
| | | 16[143] | GCCATCAAGCTCATTTTTTAACCACAAATCCATTTTCATTCTCC TATTACTACC |
| | | 12[143] | TTCTACTACGCGAGCTGAAAAGGTTACCGCGCTTTTCATTCTC CTATTACTACC |
| | | 16[175] | TATAACTAACAAAGAACGCGAGAACGCCAATTTTCATTCTCCT ATTACTACC |
| | | 14[175] | CATGTAATAGAATATAAAGTACCAAGCCGTTTTTCATTCTCCTA TTACTACC |

| | | | |
|---|---|---|---|
| | 10 nm Hexagon | 8[144] | TTATTACGAAGAACTGGCATGATTGCGAGAGGTTTTCATTCTCCTATTACTACC |
| | | 11[95] | CGAAAGACTTTGATAAGAGGTCATATTTCGCATTTTCATTCTCCTATTACTACC |
| | | 14[112] | TGTAGCCATTAAAATTCGCATTAAATGCCGGATTTTCATTCTCCTATTACTACC |
| | | 11[159] | TTCTACTACGCGAGCTGAAAAGGTTACCGCGCTTTTCATTCTCCTATTACTACC |
| | | 14[144] | GTAATAAGTTAGGCAGAGGCATTTATGATATTTTTTCATTCTCCTATTACTACC |
| | | 8[112] | TTGCTCCTTTCAAATATCGCGTTTGAGGGGGTTTTTCATTCTCCTATTACTACC |
| | 30 nm Hexagon | 14[47] | AACAAGAGGGATAAAAATTTTTAGCATAAAGCTTTTCATTCTCCTATTACTACC |
| | | 22[111] | GCCCGAGAGTCCACGCTGGTTTGCAGCTAACTTTTTCATTCTCCTATTACTACC |
| | | 6[111] | ATTACCTTTGAATAAGGCTTGCCCAAATCCGCTTTTCATTCTCCTATTACTACC |
| | | 22[175] | ACCTTGCTTGGTCAGTTGGCAAAGAGCGGATTTTCATTCTCCTATTACTACC |
| | | 6[175] | CAGCAAAAGGAAACGTCACCAATGAGCCGCTTTTCATTCTCCTATTACTACC |
| | | 14[239] | AGTATAAAGTTCAGCTAATGCAGATGTCTTTCTTTTCATTCTCCTATTACTACC |
| Maleimide | Coupling sequence | | GGTAGTAATAGGAGAATGAa[Malemid] |

**Figure S2** DNA origami design. (A) CaDNAno File and (B) Picasso Designs of the 10 nm, 5 nm and 30 nm hexagon (depicted in blue, green and red respectively). Special sequences are listed in Table (C).

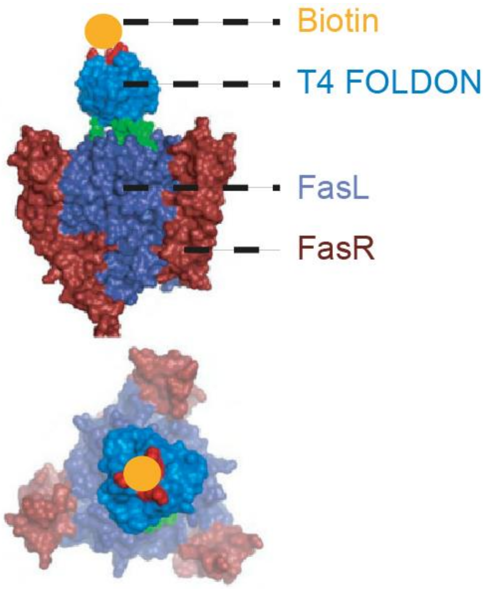


**Figure S3** Trimeric FasL with T4 FOLDON and Biotin binding to FasR. Adapted with permission from:[4].

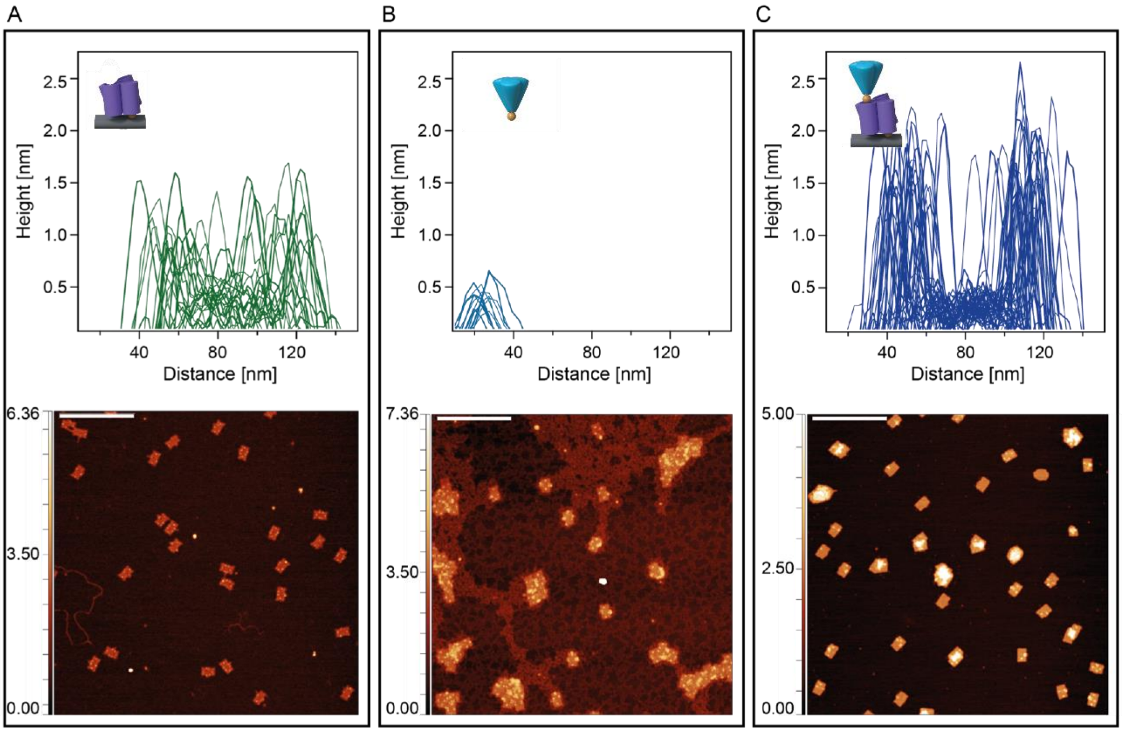


**Figure S4** Height analysis of functionalized DNA origami. AFM images of DNA origami with 30 nm hexagonal arrangements were analyzed in dry mode on AFM. (A) DNA origami only functionalized with Streptavidin. Heights were evaluated on DNA origami. Average height of Streptavidin: 1.3 ± 0.3 nm (N=35). (B) DNA origami after FasL functionalization without purification. Heights were evaluated where single FasL molecules were visible on mica. Average height of FasL: 0.5 ± 0.1 nm (N=13). (C) DNA origami after FasL functionalization and Amicon purification. Heights were evaluated on DNA origami. Average height of Streptavidin + FasL: 1.8 ± 0.3 nm (N=38, with 76% with heights above 1.5 nm).

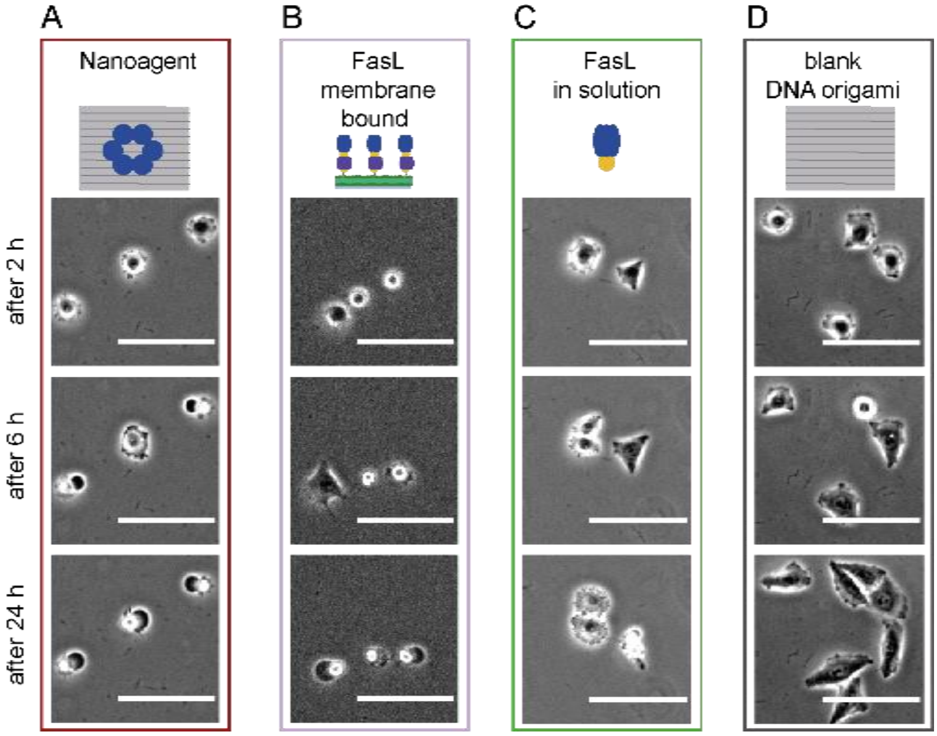

**Figure S5 Apoptosis induction depends on FasL presentation**. Representative bright field images showing cell morphologies changing in response to various FasL-functionalized surfaces after 2 h, 6 h, and 24 h: (A) FasL bound to DNA origami (red), (B) FasL bound to membrane (purple), (C) FasL in solution (green) and (D) control (grey). Cells in contact with FasL-DNA origami or FasL decorated membranes undergo apoptosis. Low number of apoptotic cells are found in the presence of FasL in solution. Cells spread and divide on controls.

**Table S6.**

| sample [units] | 1st peak [h] | 2nd peak [h] | 3rd peak [h] |
|---|---|---|---|
| Hexagon 30 nm | 14 ± 0.3 | 20 ± 2 | |
| Hexagon 10 nm | 3.1 ± 0.1 | 5 ± 1 | |
| Hexagon 5 nm | 11 ± 1 | 17 ± 1 | |
| 2-FasL 10 nm | 3.6 ± 0.1 | 7 ± 1 | |
| 2-FasL 20 nm | 3.5 ± 0.1 | 8 ± 1 | |
| FasL membrane | 6 ± 0.4 | 14 ± 1 | 21 ±1 |
| Hexagon 30 nm (mv) | 7 ± 1 | 17 ± 17 | |
| Hexagon 10 nm (mv) | 5.9 ± 0.2 | 12 ±1 | |
| Hexagon 5 nm (mv) | 7.5 ± 1 | 15 ± 0.4 | |
| 2-FasL 10 nm (mv) | 4.8 ± 0.1 | 12.5 ± 0.1 | 17.8 ± 0.3 |
| 2-FasL 20 nm (mv) | 5.5 ± 0.3 | 12 ± 0.3 | 18 ± 1 |
| Monomer (mv) | 5.6 ± 1 | 12 ±1 | 16.1 ± 0.4 |

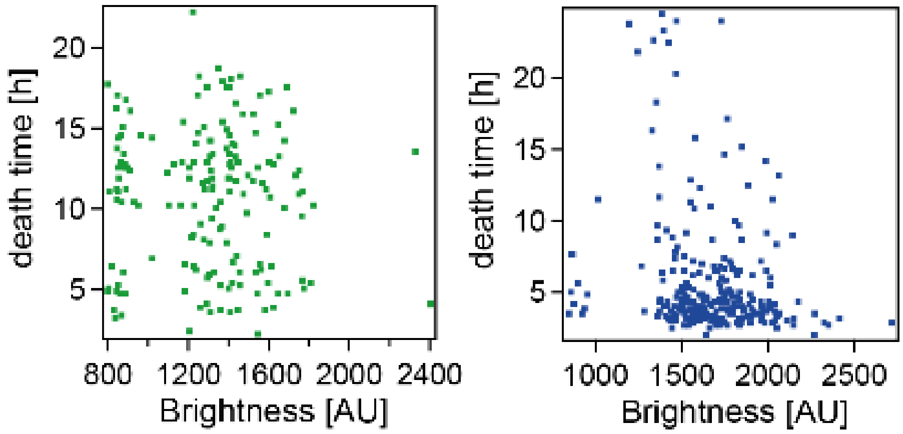


**Figure S7** HeLa cells expression of FasR-eGFP. The brightness of each cell corresponds to the amount of FasR expressed. No obvious correlation between brightness and hence receptor concentration with death time was found. Green data: DNA origamis functionalized with two FasL at a distance of 10 nm attached to SLM. Blue data: 0.5 nM hexagonal FasL-DNA origami (10 nm inter-ligand distance) in solution.

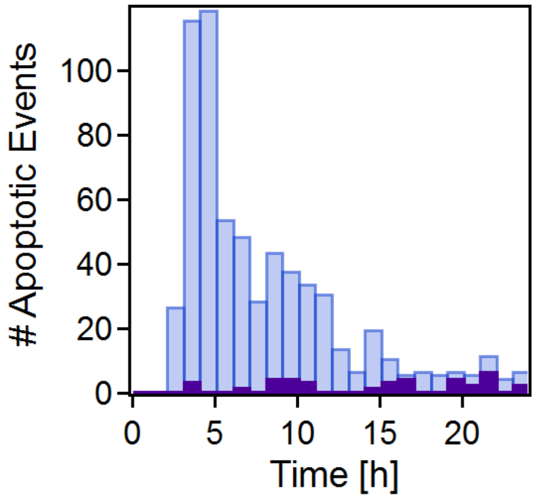


**Figure S8** Effect of cell division on time-to-death. Blue bars: all data; purple: cells dividing during measurement time.

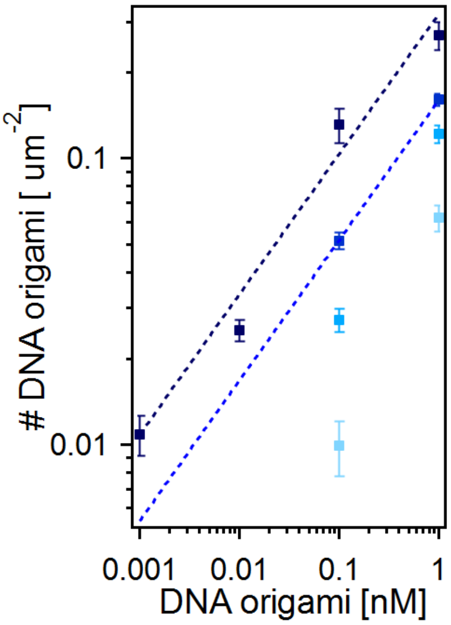


**Figure S9** Detected DNA origami per surface depending on applied concentration for different time points during measurement. To measure dose-response curves, cells were exposed to decreasing amounts of FasL-DNA origami (hexagonal arrangement with 10 nm inter-ligand spacing) by preparing surfaces at different volume concentrations of DNA origamis during incubation. The surface number concentration was calculated according to independent calibration experiments using fluorescently labeled DNA origamis and counting the number of FasL-DNA origamis per area. From dark to bright: initial number after lipid functionalization, number in cell medium, after 3 h in cell medium and after 24 h in cell medium. Fit: f(x)=a*x^b with a = 0.32 ± 0.02 $\mu m^{-2} nM^{-1}$, b = 0.49 ± 0.1 and extrapolation with b2 = b and a2 = 0.16 ± 0.001 $\mu m^{-2} nM^{-1}$.

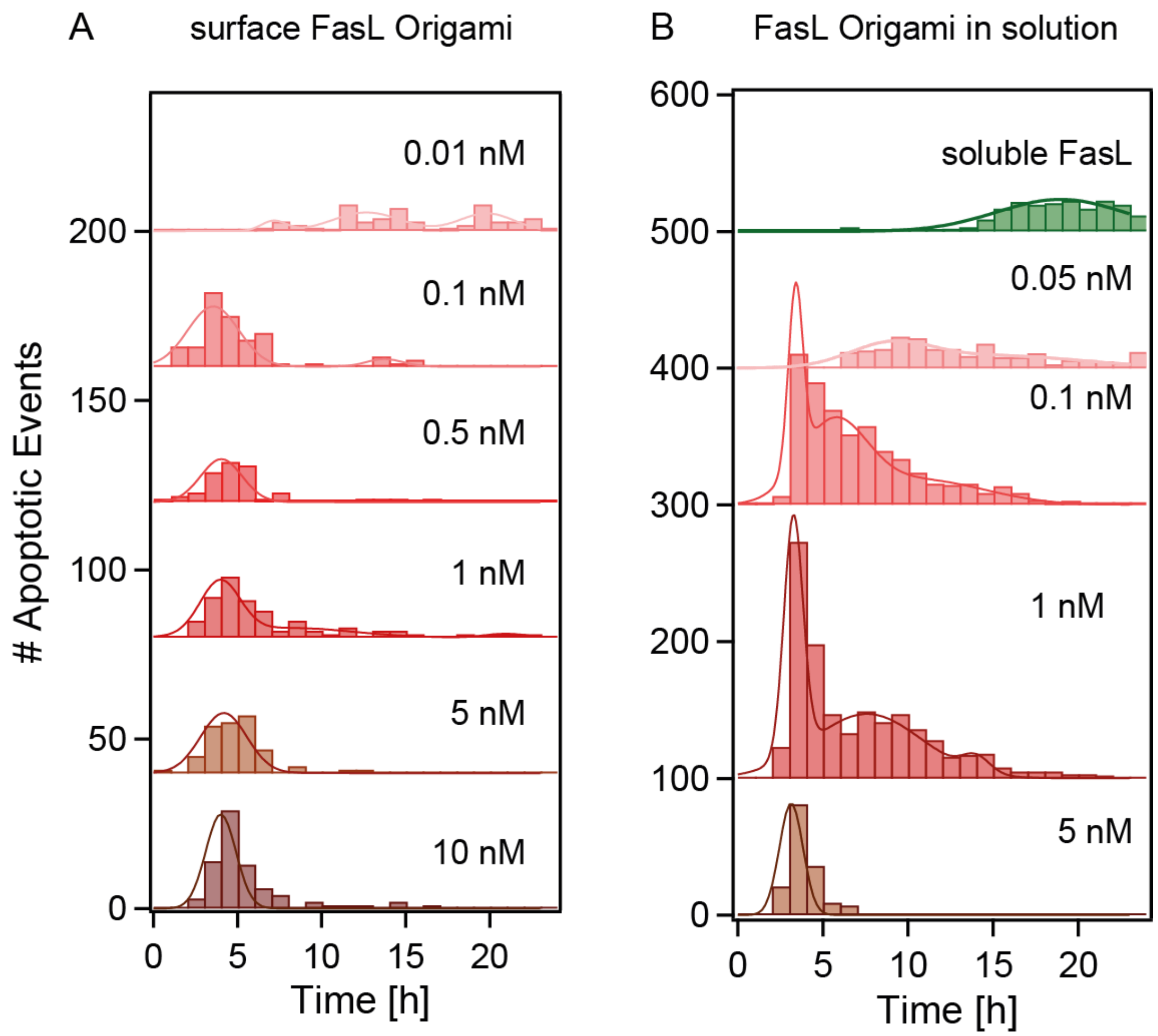


**Figure S10** Apoptotic event time histograms for FasL-DNA origami adsorbed on SLM (A) and in solution (B). For concentrations see legend.
