## Supplementary Movie Captions for "Nanoscale Organization of FasL on DNA Origami as a Versatile Platform to Tune Apoptosis Signaling in Cells"

**Movies M1** Fluorescent antibody labelling of DNA origami on SLM. (A) Before antibody labelling: only fluorescent origamis are visible (Atto488). (B) After CD178 antibody (Fluorophore: APC) addition in green and red channel fluorescence observable. 100 ms exposure and 15 frames per second.

**Movies M2** Representative 24 h timelapse movies obtained with an 10x objective for cells seeded on supported lipid membranes functionalized with FasL-DNA origami (A), or blank DNA origami (B).
